# Prediction of physical characteristics of disordered proteins using molecular simulation and physics-informed multiple machine learning strategies

**DOI:** 10.1101/2025.06.09.658593

**Authors:** Diego Linares Gonzalez, Shahana Ibrahim, Swarnadeep Seth, George Atia, Aniket Bhattacharya

## Abstract

We introduce a novel hybrid machine learning (ML) framework to predict the radius of gyration and other conformational properties of intrinsically disordered proteins (IDPs). Our model integrates sequence information with physical features derived from a coarse-grained (CG) model validated by experimental data. Specifically, we combine hidden states from sequence-based models with 23 physical features projected into a shared latent space, and apply an attention mechanism that assigns weights to each residue to highlight the most informative regions of the sequence. This attention-guided fusion significantly improves predictive accuracy across multiple metrics, including MAPE and MSE, while also enhancing confidence in the predictions. We trained and evaluated our models on Brownian dynamics (BD) simulation results for approximately 7,000 IDPs from the MobiDB database (each with *>* 99% disorder score). We find that sequence-based models consistently outperform feature-only models, with the GRU achieving the best performance among sequence-only approaches. Moreover, combining sequence and feature information further improves accuracy across all architectures, with the hybrid biGRU model delivering the best overall predictive performance. SHAP analysis reveals the relative importance of physical features, offering model explainability and guiding feature selection. Notably, using a small number of top features often reduces model complexity and improves generalization. Furthermore an integrated gradient analysis reveals that apart from the length of the IDPs, the three parameters (SCD, SHD, and *f* ^*\**^) play key role in ML predictions. Our framework provides a fast, interpretable, and scalable tool for predicting IDP behavior, enabling efficient initial screening prior to costly molecular simulations.

## I. Introduction

Over the last three decades, intrinsically disordered proteins (IDPs) have attracted growing interest [1–7]. IDPs play critical roles in numerous biological processes. Unlike folded proteins that adopt a unique three-dimensional structure to perform specific functions, IDPs adopt and evolve into many different conformations. The dynamical heterogeneities due to rapid evolution of conformational ensemble make it difficult to measure properties of IDPs experimentally [3]. The number of enlisted IDPs in various databases [8, 9] is rapidly increasing and now we know that about 40% of human proteomes are IDPs or the folded proteins have intrinsically disordered regions (IDRs) and participate in many important tasks complementing those done by the folded proteins invalidating the structure-function dogma of molecular biology. The statistical mechanics of IDP complexes have opened the door for exotic liquid-like states as IDPs form membraneless organelles through liquid-liquid and liquid-solid phase separations. The liquid-solid transition has been associated with various diseases [10, 11]. However, compared to the available IDP sequences, the availability of experimental data is far less [12]–typically carried out using small angle X-Ray scattering (SAXS) [13], single molecule Fluorescence Resonance Spectroscopy (smFRET) [14] and high field nuclear magnetic resonance (NMR) [15] methods. The highly flexible and rapidly evolving conformations of IDPs that allow them to interact with several binding partners to trigger correlated tasks, however, make the experimental studies challenging.

Machine learning (ML) algorithms have been used to partially mitigate the disparity of availability of experimental versus cataloged IDPs and IDRs (hereafter IDPs). In a previous study, we demonstrated that a simple Fully Connected Neural Network (FCNN) has the capability of predicting gyration radii with over 97% accuracy for point mutations (missense mutations) of the amino acids in an IDP [16]. Motivated by the continued success of diverse ML architectures on large datasets, we explore whether alternative architectures can further reduce the Mean Square Error (MSE). This is the central goal of the present work. A key contribution of this paper is the introduction of a **novel hybrid architecture** for predicting the radius of gyration, ⟨*R*_*g*_⟩, of IDPs. Our architecture integrates a sequence-based deep neural network (DNN), which encodes the amino acid sequence, with an FCNN that processes a set of physical features derived from coarse-grained (CG) models of IDPs validated by experimental data [17–19]. The outputs of these two branches are combined to form a unified representation for prediction. *The inclusion of physical features serves a role analogous to an attention mechanism*: it provides global biophysical context that helps the model modulate its interpretation of local sequence information. Unlike previous studies [20, 21], the hybrid design used here enhances the model’s ability to focus on sequence regions that are most informative for estimating ⟨*R*_*g*_⟩ (Fig. 3). The designed architecture is a general framework that can incorporate different sequence-based models such as recurrent neural networks (RNNs), long short-term memory networks (LSTMs), gated recurrent units (GRUs), and Transformers.

The rest of the paper is organized as follows. Section II introduces the CG model. Section III introduces IDP sequences from the MobiDB database and their characteristics based on polymer physics ideas and describes the physics-based features (input parameters) for the artificial neural net (ANN). Section IV introduces different ANN architectures and additional details of the implementation of the models. In Section V, we report the results. Finally, we summarize the key aspects of this study and provide concluding remarks in Section VI.

## II. Coarse-Grained (CG) Model for IDPs

The amino acid residues are represented as single CG beads (see Fig. 1) with interaction among 20 different beads being introduced through a 20 *×* 20 symmetric matrix first introduced by Ashbaugh-Hatch (AH) [22]. The AH potential is the result of modification of the Van der Waals interaction by adding the hydropathy terms *λ*_*ij*_ to differentiate the interaction among the amino acid beads *i* and *j*, and is given by

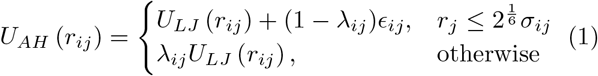

where *U*_*LJ*_ is the Lennard-Jones (LJ) potential,

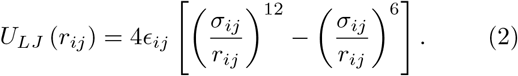

**FIG. 1.**
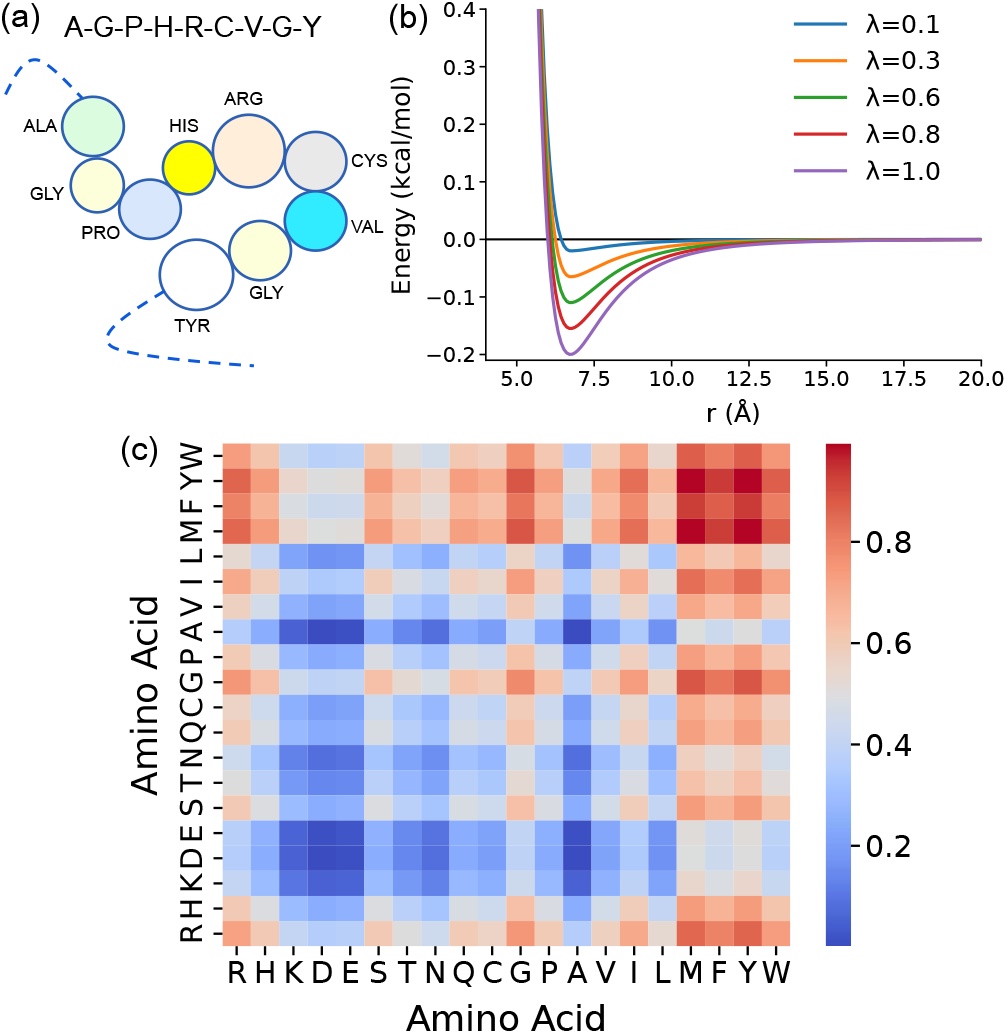
(a) CG model of single-bead IDP chain. Each bead represents an Amino acid that interacts with other beads described by the interaction potential as described in Eqns. 1–4. (b) Well depth of the AH interaction potential as a function of *λ*_*ij*_ and (c) 20 *×* 20 hydropatahy matrix as described by Eqn. 1 using CALVADOS scale [18].

Here, 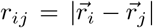 is the distance between the amino acid beads with indices *i* and *j* positioned at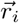 and 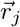 and 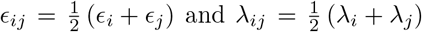 are the strength of the van der Waal interaction and average hydropathy factor between any two amino acids with indices *i* and *j*. This hydropathy factor *λ*_*ij*_ is the key ingredient of the model to differentiate interactions among IDPs.

Historically, there are many different hydropathy scales as discussed in our previous publication [19]. Here, we use the CALVADOS hydropathy scale introduced by Tesei *et al*. [18] with optimized interaction parameter *ϵ* = 0.2 kcal/mol (the model uses *ϵ*_*ij*_ = *ϵ* for all *i* and *j*) to run BD simulations to obtain the gyration radii (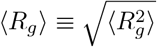for brevity) of approximately 7000 IDPs to train several neural network architectures as discussed in section IV. *ϵ* = 0.2 in this model produce near identical results (MSE) with the hydropathy scale due to Dignon [17] with *ϵ* = 0.1 as shown in our recent publication [19].

We introduce a harmonic bond potential with spring constant *k*_*b*_ = 8033 kJ/(mol·nm^2^) = 1920 kcal/(mol·nm^2^ (Eqn. 3)

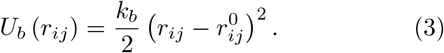

between two consecutive amino acid residues *i* and *j* = *i ±* 1 with the equilibrium bond length 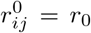. Here 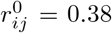 is the distance between *α*-Carbon atoms for the successive amino acids. Thus, we exclude the EV interaction among the bonded neighbors.

The charged species of the amino acids interact with Screened-Coulomb (SC) interaction (Eqn. 4) given by

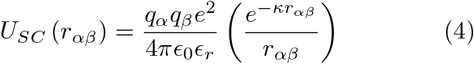

where the indices *α* and *β* refer to the subset of the indices *i* and *j* for the charged amino acids, *ϵ*_*r*_ is the dielectric constant of water, and *κ* is the inverse Debye screening length [23]. The inverse Debye length *κ*^−1^ is dependent on the ionic concentration (I) and expressed as

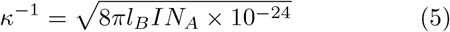

where *N*_*A*_ is the Avogadro’s number and *l*_*B*_ is the Bjerrum length,

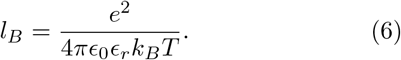

Our model slightly differs from that of Tesei et al. and Dignon et al. in that we introduce a temperature-dependent dielectric constant of water as expressed by the empirical relation [24]

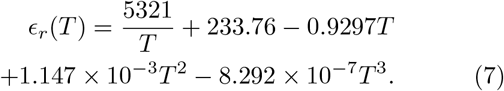

This accounts for the typical decrease of the dielectric constant at higher temperature. Without this term the the electrostatic interactions may be overestimated leading to unrealistic protein conformations. In addition to introducing this temperature dependent architecture, we keep the bare charge of the amino acid at two ends the same. Previously, we called this model as HPS2 model and the model due to Dignon with similar changes as HPS1 model [19]. In this paper, to train an ANN, we use physical characteristics drawn from the HPS2 scale.

## III. Physical Characteristics of IDPs and Physics-based Features to train ANN

We gathered approximately 7000 IDP FASTA sequences from the MobiDB database with the predicted disordered score exceeding 99%. The number of amino acids in the sequence *N* (actual length *L* = 0.38*N* nm), ranges between 20 − 150 with almost 2300 IDPs of length between 40 to 60 as shown in Fig. 2(a). The IDPs are well known for containing a larger fraction of disorder promoting residues such as Lysine, Glutamine, Serine, Proline, Glycine, Arginine, and Glutamic acid. From the dataset of 7000 IDPs, we calculate the propensity of occurrence of individual amino acids and their relative probability. To train ML models to predict ⟨*R*_*g*_⟩, we use a training/validation split of 0.8:0.2, with a separate dataset of 31 IDPs reserved for testing. IDPs are mostly polyampholytes and polyelectrolytes. In addition, each amino acid bead has different mass and hydropathy index. Therefore, it is expected that the length of the chain and the distribution of the positive and negative charge resides as well as their hydropathy values will affect the conformational properties. We have used characteristics that depend on the sequence length, masses, charges and hydropathy indices of the amino acids and some distance dependent quantities, such as SCD and SHD to characterize a given IDP. The input parameters are introduced in Table I. Further details of the input parameters are provided in the supplementary section.

**TABLE I.** Feature descriptions and notations.

| Notation | Feature Description |
| --- | --- |
| $N$ | Length |
| $S_{IDP}^{all}$ | Sequence Entropy |
| $q_{net}$ | Charge per Unit Length |
| $N_q^{+1}$ | Positive Charge Count |
| $N_q^{+1/2}$ | Half Positive Charge Count |
| $N_q^{-1}$ | Negative Charge Count |
| $N_q^0$ | Neutral Charge Count |
| $S_{IDP}^q$ | Charge Entropy |
| $\langle \lambda \rangle$ | Avg Hydropathy |
| SHD | Sequence Hydropathy Decoration |
| SCD | Sequence Charge Decoration |
| $f^*$ | $f^*$ |
| $M_{CM}$ | Center of Mass |
| $Q_{CM}$ | Center of Charge |
| $\lambda_{CM}$ | Center of Hydropathy |
| $\delta M$ | MF STD Mass |
| $\delta Q$ | MF STD Charge |
| $\delta \lambda$ | MF STD Hydropathy |
| SBCS | SBCS |
| $l_{patch}^{(-1)}$ | Patch Neg1 |
| $l_{patch}^{(+1)}$ | Patch POS1 |
| $l_{patch}^{(+1/2)}$ | Patch Pos half |
| $l_{patch}^{(0)}$ | Patch Zero |

**FIG. 2.**
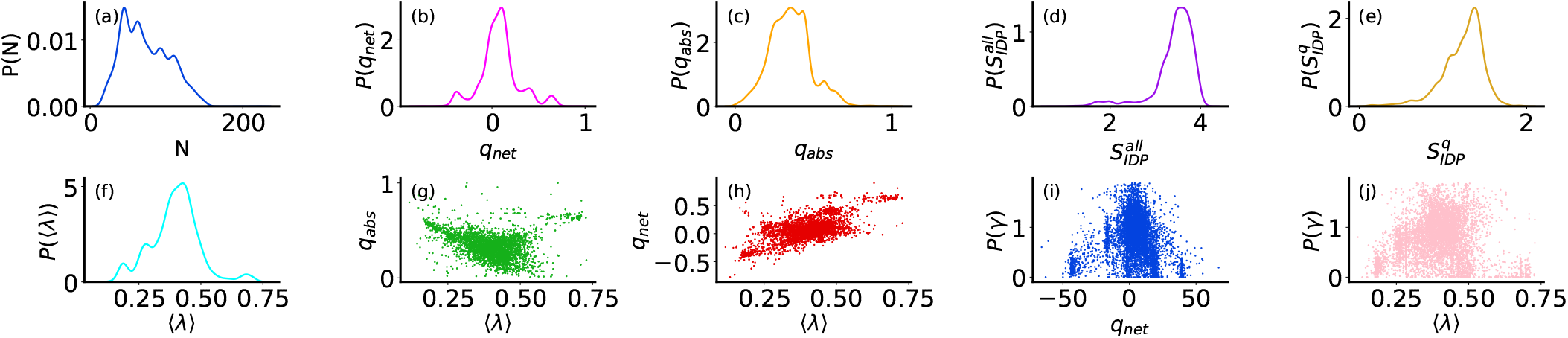
(a) Physical characteristics of 6500 IDPs with a disorder predictor score of ≥ 99%, sourced from the MobiDB database. (a)-(f) the distribution of IDP lengths, Net Charge Per Residue (NCPR) *q*_*net*_, Absolute Charge Per Residue (ACPR) *q*_*abs*_, sequence entropy 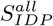, charge entropy 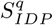, and average hydropathy ⟨*λ*⟩. (g)-(h) depicts ACPR and NCPR as a function of⟨*λ*⟩. (i)-(j) exhibits skewness 𝒮 as a function of *q*_net_ and ⟨*λ*⟩.

## IV. Neural Network Architectures

In this section, we describe the ML architectures used in this study, including a novel hybrid architecture that integrates sequence information with physical features. We compare against sequence-only models based on recurrent neural networks (RNN, LSTM, GRU, biLSTM, biGRU) and Transformer-based architectures, as well as a feature-only model that employs an FCNN using the set of physical features described in Section III.

### A. Hybrid Architecture

Our proposed hybrid architecture integrates sequential data processing with physical feature incorporation to enhance predictive performance for IDP analysis. In this architecture, illustrated in Fig. 3, we combine sequence-based models—RNN, LSTM, biLSTM, GRU, and biGRU—with a per-residue fusion mechanism and an attention-based aggregation strategy. This framework can handle variable-length sequences, and using their associated physical features offers accurate outputs.

**FIG. 3.**
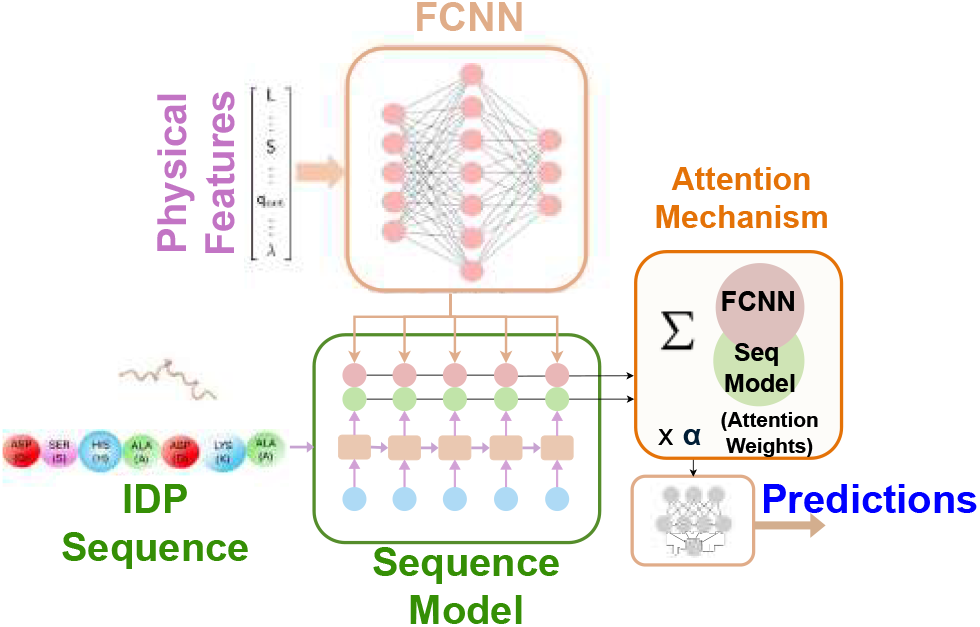
Hybrid model combining sequences and physical features. Hidden states from sequence models (LSTM, GRU, RNN variants) are concatenated with physical features mapped to a 32-dimensional space. An attention mechanism assigns weights to each residue, emphasizing informative regions. The weighted sum is passed to a final output layer for prediction.

A key novelty lies in the residue-wise fusion of the sequence and feature inputs, combined with an attention mechanism that gives more importance to critical residues, helping the model adaptively focus on different parts of the sequences. The main components of our hybrid architecture are detailed as follows:

#### Sequence processing

The input sequence, representing IDP residues, is processed using one of five the recurrent architectures: RNN, LSTM, biLSTM, GRU, or biGRU. Each residue in a sequence is represented using one-hot encoding over the 20 standard amino acids. Hence, each IDP sequence is represented as an input matrix 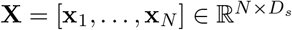, where **x**_*n*_ is the one-hot encoding corresponding to the *n*th residue, *N* is the sequence length and *D*_*s*_ = 20 is the total number of possible residues. For each sequence, the recurrent layer produces hidden states **H** = [**h**_1_, …, **h**_*N*_] ∈ ℝ^*H×N*^, where *H* is the dimension of each vector **h**_*n*_. Unidirectional models (RNN, LSTM, GRU) output **h**_*n*_, while bidirectional variants (biLSTM, biGRU) combine forward and backward passes to yield **h**_*n*_ ∈ ℝ^2*H*^.

LSTM models additionally maintain a cell state **c**_*n*_ to capture long-range dependencies. GRUs, in contrast, use a gating mechanism to update hidden states directly. All variants follow the general update form:

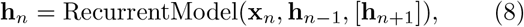

with backward context **h**_*n*+1_ used only in bidirectional models.

#### Physical feature processing

The physical features are processed through a fully connected layer as follows:

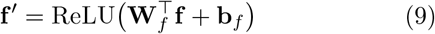

where 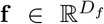 is the input feature vector with *D*_*f*_ denoting the total number of physical features. Here 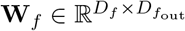 and 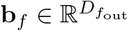 are the weight matrix and bias, respectively, ReLU(*x*) := max(0, *x*) is the activation function and 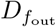 denotes the dimension of the feature embedding vector **f** ^*′*^. Here, *D*_*f*_ = 23, i.e., there are 23 input features, and we fix 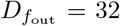. This step prepares the features to be fused with the hidden states in following steps.

#### Residue-wise fusion

An important step of our hybrid architecture is the residue-wise fusion of sequence and feature information. For each residue *n*, the hidden state **h**_*n*_ is concatenated with the processed features **f** ^*′*^:

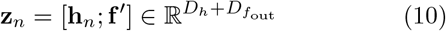

where *D*_*h*_ = *H* for unidirectional models and *D*_*h*_ = 2*H* for bidirectional models. The fused representation **z**_*n*_ is passed through another fully connected layer to produce a refined representation:

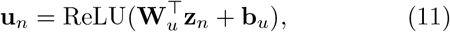

where 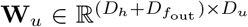 and 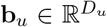 are the fusion layer parameters, and *D*_*u*_ = 128 is the fusion dimension. The main goal of this per-residue fusion is to integrate sequence information with global physical features at each residue, allowing the model to capture complex interactions between sequence and feature data.

#### Attention mechanism

To aggregate the variable-length fused representations 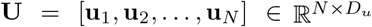 into a fixed-length vector, we employ an attention mechanism that learns different weights for each position. Here, *N* represents the length of each input sequence and *D*_*u*_ is the dimension of each vector **u**_*n*_. Since sequences can vary in length, a padding mask is applied to exclude the padded positions from the calculations. The attention scores are computed as follows:

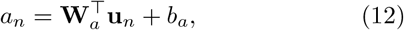

where 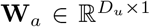 and *b*_*a*_ ∈ ℝ are the weight matrix and the bias term, respectively, producing a score *a*_*n*_ for each residue position *n* = 1, …, *N*.

The attention scores are further normalized using a softmax function to produce the final attention weights as follows:

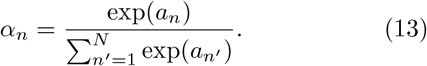

Using the attention weights, the final sequence-feature embedding vector is computed as a weighted sum of the fused representations:

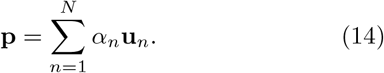

This fixed-length vector 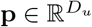 is then passed through a final two-layer fully connected network to produce the output prediction **y**:

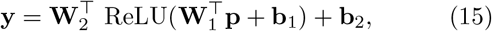

where 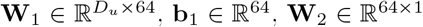, and **b**_2_ ∈ ℝ denote the weight matrices and bias terms of the two-layer output head, respectively.

In the Transformer-based hybrid model, we average the encoder outputs across the sequence to obtain a fixed-length summary representation. We then concatenate this with the processed numeric features and pass the combined vector through fully connected layers to generate the final prediction.

### B. Sequence-Only Architecture

We compare the performance of our proposed hybrid model against different sequence-only architectures. Sequence-based architectures such as RNN, LSTM, GRU, and Transformers are designed to capture the order and temporal dependencies within sequences.To process the sequences, we use one-hot encoding, representing each amino acid residue as a binary vector of length 20. In our implementation, the RNN, GRU, and LSTM models each use two layers with a hidden size of 64. The GRU and LSTM models are also implemented in a bidirectional configuration, with forward and backward hidden states concatenated before the output layer. For all sequence models, input sequences were packed according to their lengths, and initial hidden states were zero-initialized. In the Transformer model, we project 20-dimensional amino acid inputs into a 64-dimensional embedding space and add randomly initialized, learnable positional encodings. We use two encoder layers, each with four attention heads, a feedforward dimension of 256, and a dropout rate of 10%. To handle sequences of varying lengths, we apply a padding mask during attention computation. The encoder outputs are mean-pooled across the sequence and passed through a linear layer to produce the final prediction.

### C. Feature-Only Architecture

We also consider an FCNN that operates solely on physical (non-sequential) features. The FCNN model consists of four fully connected layers: an input layer mapping to 128 dimensions, followed by two hidden layers with 64 and 32 units, and a final output layer. Each layer uses the rectified linear unit (ReLU) activation function, which introduces nonlinearity and helps mitigate the vanishing gradient problem common in deep networks [25]. To reduce the risk of overfitting—a common phenomenon in machine learning where the model learns to memorize training data rather than generalizing to unseen samples—we apply dropout [26] after the first layer at a rate of 10%, which is a regularization technique that randomly sets a fraction of the hidden unit outputs to zero during training to improve the model’s generalization performance.

### D. Other Implementation Details

All models were implemented in PyTorch and trained using the Adam optimizer with a learning rate of 0.001. The Mean Absolute Error (MAE) was used as the loss function. Training was performed for up to 650 epochs with a batch size of 32, using early stopping with a patience of 60 epochs—that is, training stops if validation performance does not improve for 60 consecutive epochs. Each experiment was repeated across 3 random trials. To improve training stability, inputs involving the radius of gyration ⟨*R*_*g*_⟩ were normalized by dividing ⟨*R*_*g*_⟩ by 0.38*L*^0.56^, where *L* denotes the sequence length. This scaling relationship between ⟨*R*_*g*_⟩ and *L* was introduced by Seth et al. [19] and validated through extensive testing on a dataset of 2000 simulated proteins, as well as 31 well-studied simulated IDPs. By incorporating this normalization, we prevent the model from overfitting to this dominant scaling law and instead encourage it to discover additional meaningful patterns in the data. Normalizing ⟨*R*_*g*_⟩ in this way leads to more balanced learning and ultimately improves prediction accuracy. All numerical features, excluding length, and ⟨*R*_*g*_⟩, were normalized using the Python preprocessing library `StandardScaler`.

### E. Evaluation Metrics

To evaluate the performance of the machine learning predictions on test data, we use different metrics such as mean squared error (MSE), mean absolute error (MAE), and mean absolute percentage error (MAPE), which are defined as follows:

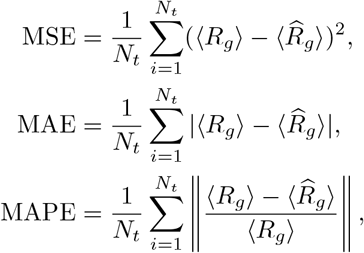

where 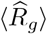 is the predicted ⟨*R*_*g*_⟩ by the model and *N*_*t*_ is the total number of test sequences.

## V. Results

We evaluate the predictive performance of various models using the MobiDB dataset. Model selection was based on validation performance, with the best results summarized in Table II and in Figure 4, where we present the average MAPE results on the test data for various models. The figure indicates that the sequence-based models outperformed feature-only models on the test data, with the GRU model achieving the best performance among sequence-only approaches (MAPE: 2.27%). In comparison, the FCNN model, which relies solely on physical features, delivered a competitive performance (MAPE: 3.18%), despite having fewer parameters. The FCNN model is computationally less expensive and faster to train. Moreover, the feature-based FCNN model offers the advantage of explainability, allowing predictions to be interpreted using feature importance analysis techniques such as SHAP [27] (see Fig. 7). Finally, the proposed Hybrid biGRU model, which combines sequence-based and feature-based information, achieved the best predictive performance among all tested architectures. In particular, as shown in Table III, it reduced the MAPE from 3.18% *±* 0.47% with features-only input and 2.27% *±* 0.19% with the best sequence-only model (GRU), to 1.98% *±* 0.09% when combining both modalities. This corresponds to a relative improvement of approximately 37.7% over the features-only model and 12.7% over the best sequence-only model. The consistent performance gains across trials underscore the value of integrating physical features with sequence data, as the additional information enhances the model’s ability to generalize. This trend is further illustrated in Figure 5, which provide a detailed comparison between the predicted and simulated *R*_*g*_ values for the 31 IDPs using the Hybrid biGRU and the FCNN model. One can note in Figure 5 that the proposed Hybrid biGRU model demonstrates superior accuracy, with a lower mean squared error (MSE = 0.01) and a higher coefficient of determination (*R*^2^ = 0.98), compared to the FCNN model (MSE = 0.03, *R*^2^ = 0.96). Moreover, we observe that both models exhibit larger predictions errors for certain proteins, such as OPN and ProTan. We also present the specific ⟨*R*_*g*_⟩ predictions in Table IV.

**TABLE II.**
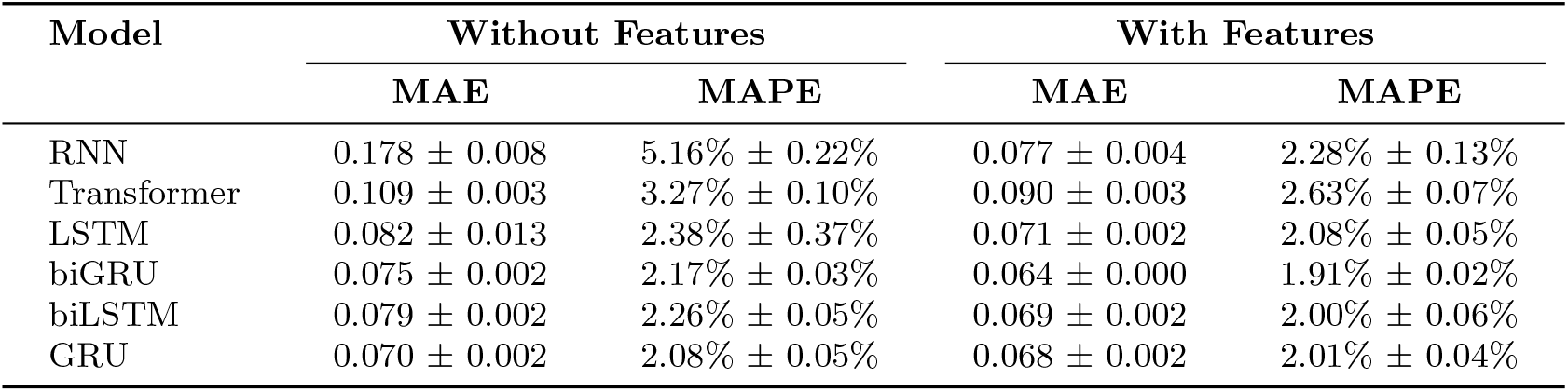
Prediction performance *on the validation dataset* for various sequence-based architectures, with and without additional features, trained and tested on MobiDB [8] sequences with disorder scores greater than 99%.

**TABLE III.** Prediction performance on the test dataset for various sequence-based architectures, with and without additional features, trained and tested on MobiDB [8] sequences with disorder scores greater than 99%.

| Model | Without Features |  | With Features |  | With Top 5 Features |  |
| --- | --- | --- | --- | --- | --- | --- |
|  | MAE | MAPE | MAE | MAPE | MAE | MAPE |
| RNN | $0.196 \pm 0.034$ | $5.60\% \pm 0.91\%$ | $0.094 \pm 0.010$ | $2.67\% \pm 0.36\%$ | $0.082 \pm 0.005$ | $2.26\% \pm 0.14\%$ |
| Transformer | $0.098 \pm 0.010$ | $2.71\% \pm 0.28\%$ | $0.099 \pm 0.010$ | $2.79\% \pm 0.26\%$ | $0.096 \pm 0.013$ | $2.66\% \pm 0.41\%$ |
| LSTM | $0.086 \pm 0.011$ | $2.43\% \pm 0.39\%$ | $0.080 \pm 0.007$ | $2.25\% \pm 0.15\%$ | $0.080 \pm 0.004$ | $2.25\% \pm 0.10\%$ |
| biGRU | $0.086 \pm 0.012$ | $2.35\% \pm 0.33\%$ | $0.070 \pm 0.003$ | $1.98\% \pm 0.09\%$ | $0.073 \pm 0.004$ | $2.05\% \pm 0.10\%$ |
| biLSTM | $0.083 \pm 0.006$ | $2.31\% \pm 0.14\%$ | $0.078 \pm 0.001$ | $2.17\% \pm 0.03\%$ | $0.078 \pm 0.006$ | $2.18\% \pm 0.14\%$ |
| GRU | $0.082 \pm 0.005$ | $2.27\% \pm 0.19\%$ | $0.079 \pm 0.006$ | $2.18\% \pm 0.14\%$ | $0.077 \pm 0.002$ | $2.12\% \pm 0.09\%$ |

**TABLE IV.** Comparison of the predicted ⟨*R*_*g*_⟩ between the Simulation (5-th Column) and (i) the Hybrid biGRU Model (3rd column) and the FCNN model (4th Column). We compared our results with 31 IDPs from the test dataset. The last two columns indicate the corresponding percentage error between the prediction and the simulation.

| IDP name | $N$ | $\langle R_g^{hGR} \rangle$ | $\langle R_g^{FCN} \rangle$ | $\langle R_g^{Sim} \rangle$ | $\langle \delta R_g^{hGR} \rangle$ | $\langle \delta R_g^{FCN} \rangle$ |
| --- | --- | --- | --- | --- | --- | --- |
| HST5 | 24 | 1.42 | 1.36 | 1.40 | 1.58 | -2.65 |
| IN | 60 | 2.25 | 2.25 | 2.20 | 2.04 | 1.91 |
| L | 64 | 2.33 | 2.37 | 2.34 | -0.43 | 1.27 |
| CspTm | 67 | 2.26 | 2.25 | 2.22 | 1.53 | 1.24 |
| ACTR | 71 | 2.53 | 2.59 | 2.56 | -1.09 | 1.05 |
| Nuclp153 | 81 | 2.45 | 2.47 | 2.42 | 0.87 | 1.83 |
| SH4UD | 85 | 2.66 | 2.65 | 2.66 | -0.23 | -0.28 |
| SIC1 | 90 | 2.85 | 2.82 | 2.82 | 0.99 | -0.11 |
| p53 | 93 | 3.04 | 3.14 | 3.05 | -0.36 | 2.74 |
| CoINT | 98 | 2.82 | 2.72 | 2.72 | 3.83 | 0.33 |
| K19 | 99 | 3.03 | 2.90 | 3.07 | -1.24 | -5.45 |
| R17 | 100 | 2.97 | 2.86 | 3.08 | -3.38 | -6.95 |
| p15PAF | 111 | 3.16 | 3.13 | 3.16 | 0.13 | -0.91 |
| ProTaN | 112 | 4.29 | 4.26 | 4.67 | -8.24 | -8.89 |
| R15 | 114 | 3.12 | 3.23 | 3.15 | -0.83 | 2.72 |
| hCyp | 115 | 3.00 | 3.01 | 2.95 | 1.66 | 1.91 |
| ERMTADn | 122 | 3.13 | 3.12 | 3.21 | -2.52 | -2.97 |
| ProTaC | 129 | 4.26 | 3.93 | 4.27 | -0.30 | -7.94 |
| K18 | 130 | 3.53 | 3.41 | 3.45 | 2.47 | -1.01 |
| hNHE1cdt | 131 | 3.41 | 3.31 | 3.49 | -2.21 | -4.90 |
| sNase | 136 | 3.57 | 3.71 | 3.68 | -2.93 | 0.64 |
| synuclein | 140 | 3.62 | 3.41 | 3.79 | -4.54 | -10.02 |
| FhuA | 143 | 3.51 | 3.50 | 3.48 | 0.69 | 0.43 |
| K17 | 145 | 3.79 | 3.65 | 3.71 | 2.16 | -1.60 |
| K27 | 167 | 4.12 | 3.93 | 4.17 | -1.17 | -5.86 |
| K10 | 168 | 3.94 | 3.81 | 3.94 | 0.18 | -3.22 |
| K16 | 176 | 4.23 | 4.05 | 4.23 | 0.05 | -4.27 |
| An16 | 185 | 4.00 | 4.06 | 3.72 | 4.78 | 9.02 |
| K25 | 185 | 4.02 | 4.01 | 4.01 | 0.22 | -0.13 |
| K32 | 198 | 4.56 | 4.29 | 4.61 | -1.21 | -7.06 |
| OPN | 219 | 4.65 | 4.70 | 4.44 | 4.91 | 5.89 |

**FIG. 4.**
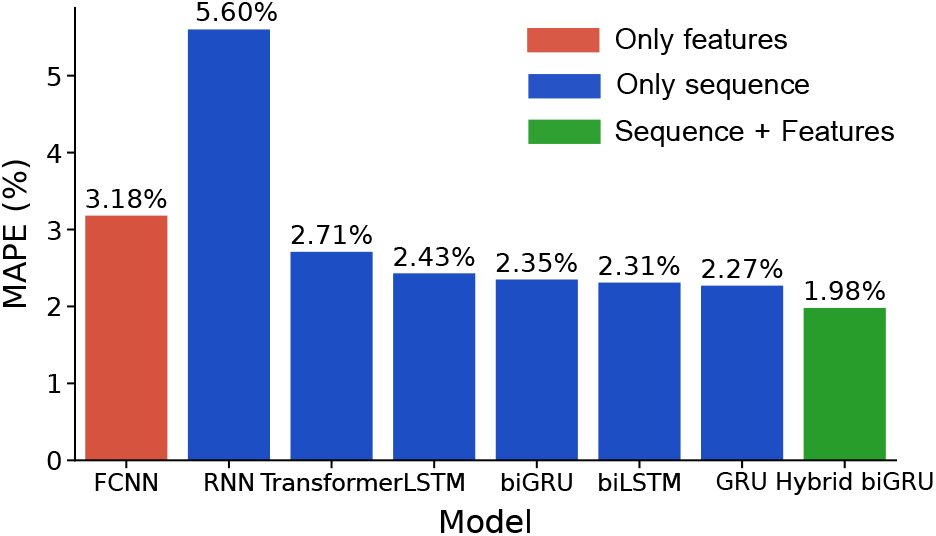
Average MAPE results on the test dataset for ⟨*R*_*g*_⟩ prediction by different AI models using (i) features only (FCNN), (ii) using sequences only (RNN, Transformer, LSTM, biLSTM, biGRU and GRU), and finally (iii) a hybrid method (Hybrid biGRU) using both sequence and features which gives least error.

**FIG. 5.**
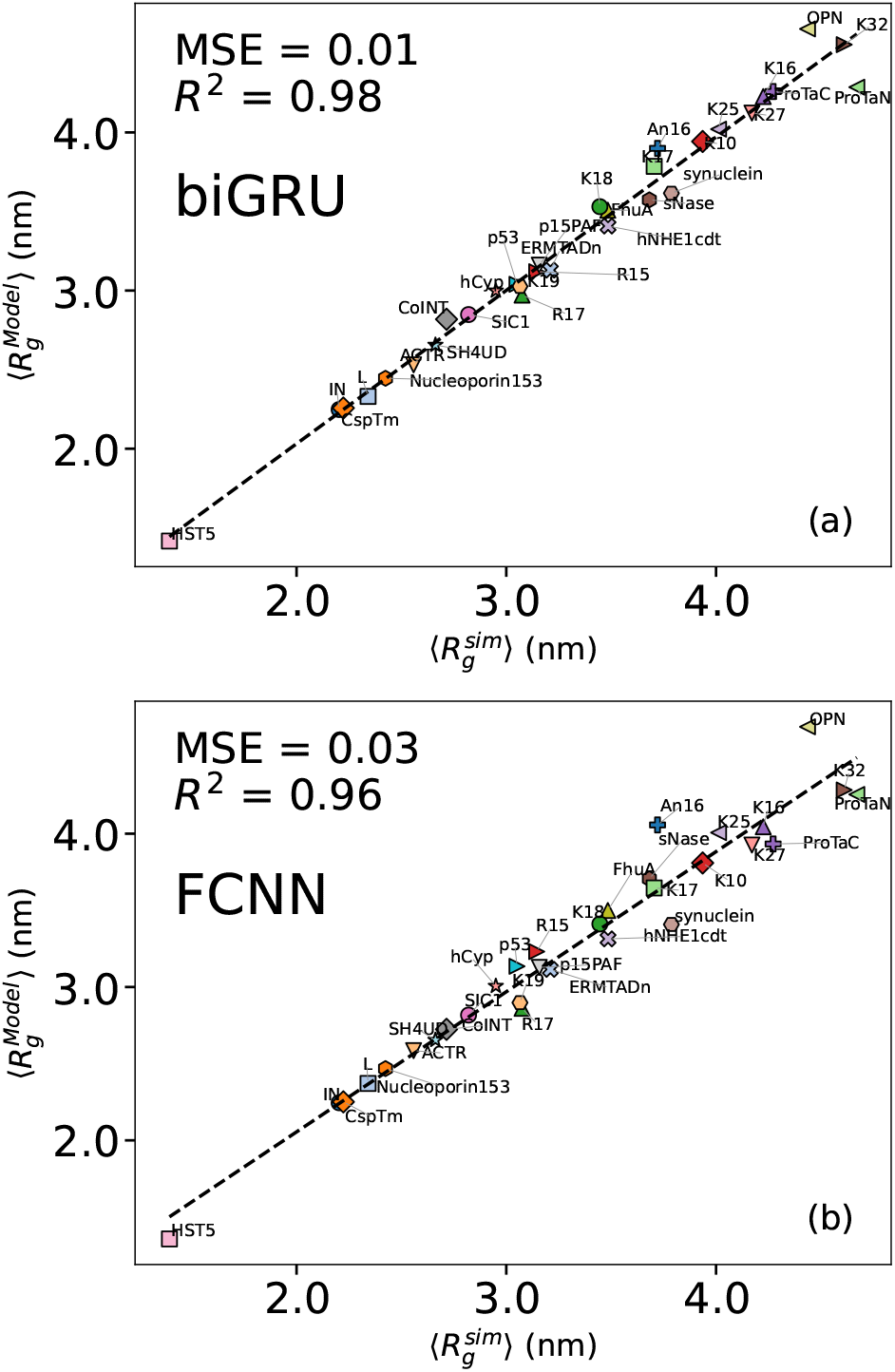
(a) Predictions for each of the 31 IDPs using the (a) Hybrid biGRU model. (b) the FCNN model

**FIG. 6.**
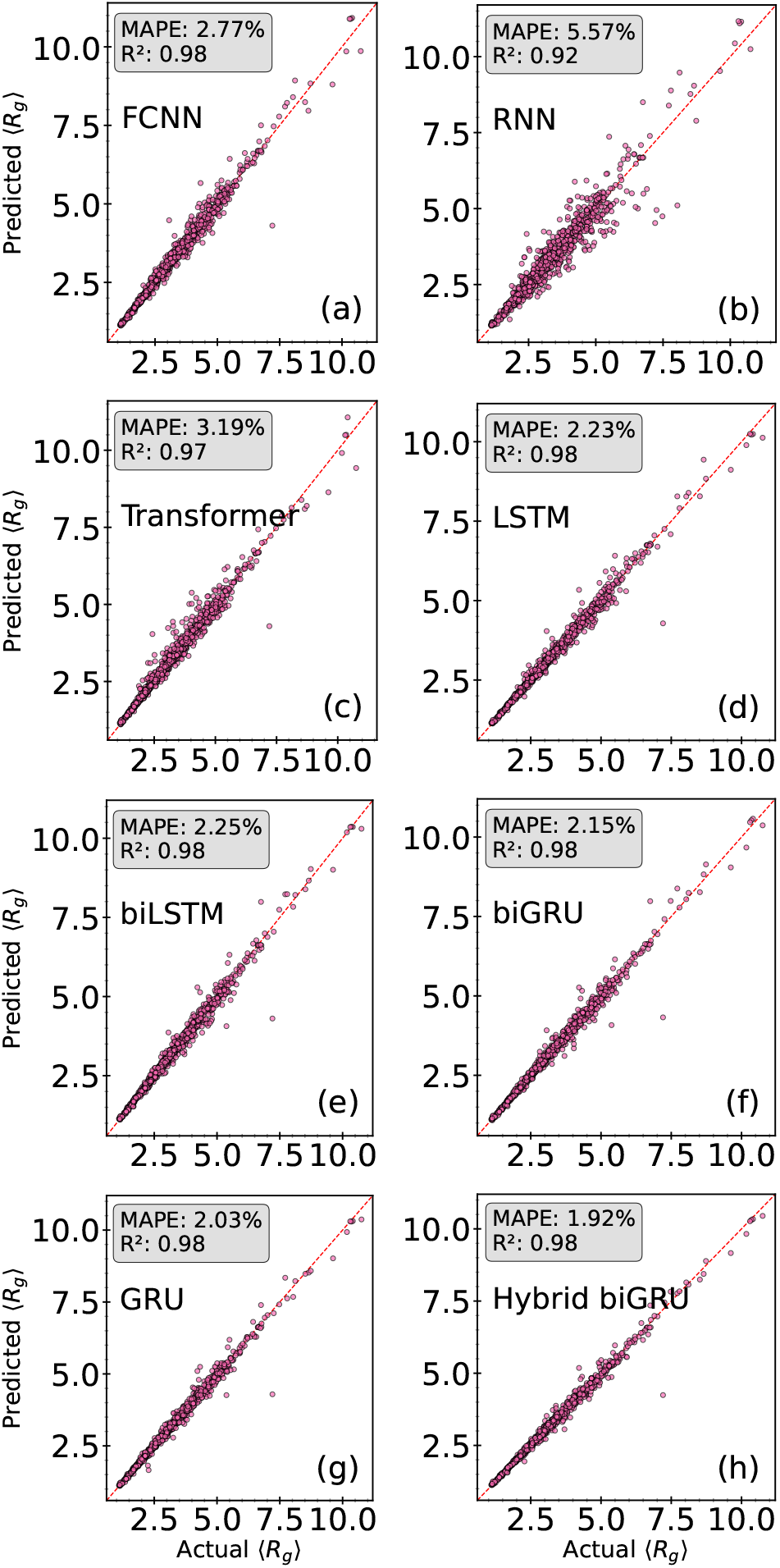
Comparison of predicted vs actual ⟨*R*_*g*_⟩ values on the validation dataset across different models: GRU (only sequence), biGRU (only sequence), Hybrid GRU (sequence + features), FCNN (only sequence), RNN (only sequence), and Transformer (only sequence). The predictions were made on protein sequences from the MobiDB [8] database with disorder scores above 99%. The plots show the mean absolute percentage error (MAPE) and the coefficient of determination (*R*^2^), which indicates the proportion of variance in the actual values explained by the model, with values closer to 1 reflecting better predictive performance.

**FIG. 7.**
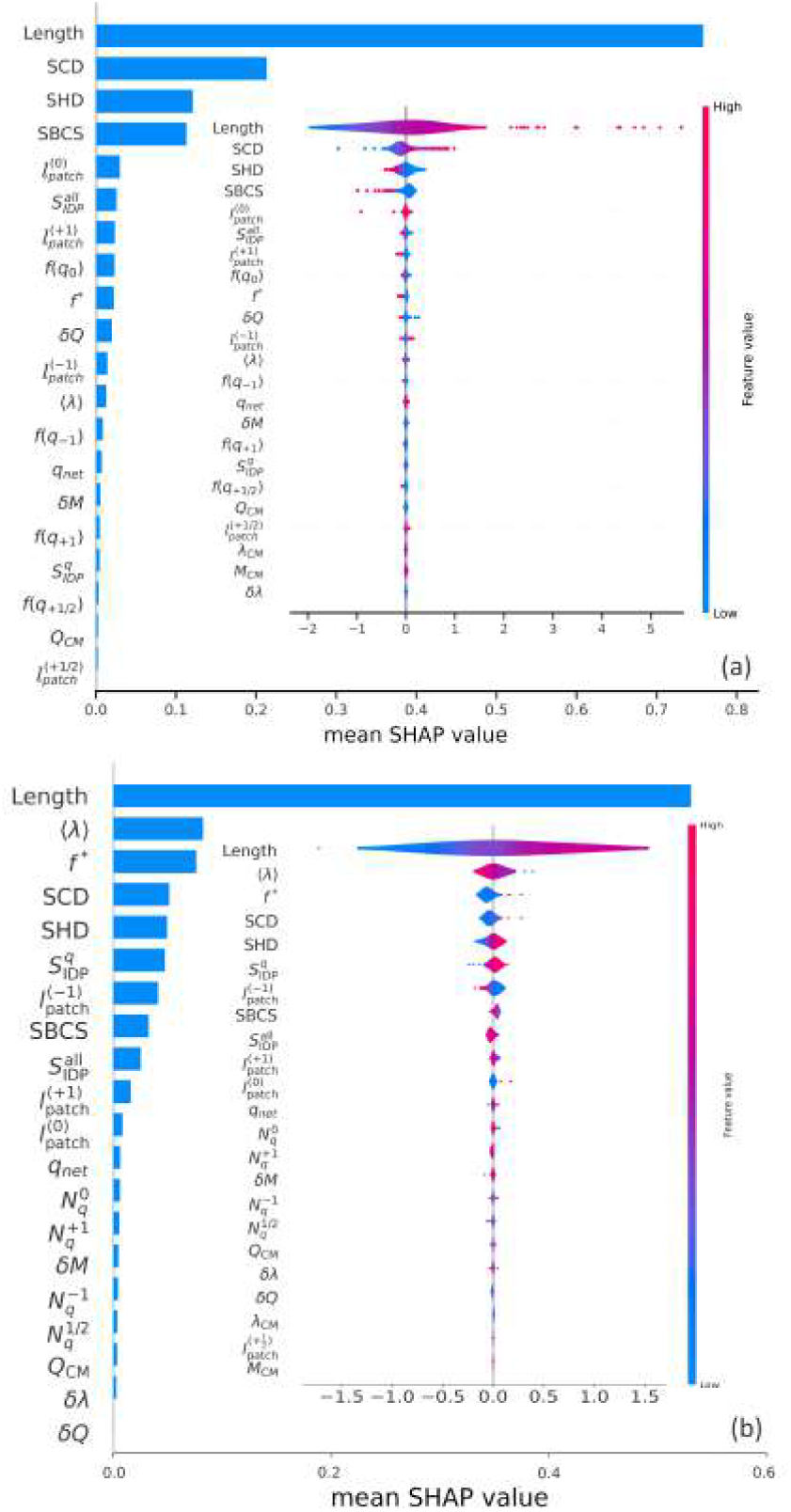
(a) SHAP summary from Seth et al (2025) [16]. Without ⟨*R*_*g*_ ⟩ normalization. (b) SHAP summary of our proposed FCCN model. Without ⟨*R*_*g*_ ⟩ normalization

In Table III, we conducted an ablation study of the hybrid model where we incorporate features into different sequence architectures. *A key observation is that our hybrid framework consistently outperforms the sequence-only variants across different model architectures*. This reinforces our hypothesis that adding physics-based features guides the learning process and improves overall outcomes.

### Explainability and feature importance

In this section, we evaluate our model’s explainability by identifying the key features that influence its predictions of ⟨*R*_*g*_⟩.

We evaluate model performance using only the top five most important features—identified via SHAP [28] and Integrated Gradients feature attribution [29] methods— in combination with the sequence data. The selected features were Length, ⟨ *λ*⟩, SHD, SCD, and *f* ^*\**^. In this setting, the models were retrained from scratch using just these selected features. Interestingly, the results showed that a reduced feature set can yield similar or even better performance. For example, the GRU model achieved a MAPE of 2.12%, compared to 2.18% when using all features. This suggests that removing less informative or redundant features can improve generalization, reduce overfitting, and lead to more efficient models without sacrificing accuracy.

In Fig. 7, we present a comparative SHAP analysis between the results reported in [16] and those obtained in this work. To enable a fairer comparison, we removed the ⟨*R*_*g*_⟩ normalization from our method to align more closely with the settings used in [16]. While Length appears to be a critical feature in both models, the relative importance of the remaining variables differs significantly, largely due to differences in model architecture. In [16], the FCNN model is composed of eight hidden layers with alternating ReLU and tanh activation functions.

In the supplementary section, we present a thorough SHAP analysis for each model that incorporates feature inputs. Since gyration radii by and large follow the scaling relation for the homopolymer ⟨*R*_*g*_⟩ ~ *L*^*ν*^ with *ν* ∼ 0.5 − 0.6 [19], it is expected that Length will remain the most dominant contributing factor. This is consistent with this analysis showing that Length remains the dominant variable across all hybrid models, while the importance of other variables slightly diminishes—suggesting that the sequence-based models likely learn these relationships across variables.

We also performed an *Integrated Gradients* analysis for each model with further details and figure in the supplementary section. This method was introduced by Sundararajan *et al*. [29], aiming to measure how a neural network’s output changes by tracking gradients from a baseline input. The analysis revealed that the top three most important features across all models were consistently Length, *f* ^*\**^, and SCD. Notably, SHD appeared as the fourth most important feature in 3 out of the 7 evaluated models.

## VI. Summary and Outlook

In conclusion, our results demonstrate that ML algorithms, particularly those that integrate both sequence information and physical features, can effectively predict conformational properties of IDPs. Through extensive evaluation across several architectures and interpretability analysis using SHAP values, we find that hybrid models consistently outperform sequence-only and feature-only baselines, with the biGRU-based hybrid achieving the best predictive performance. This supports our central hypothesis that combining biophysically motivated features with learned sequence representations improves both accuracy and confidence in predictions.

We also observe that the attention mechanism in the hybrid models provides interpretability. Furthermore, we find that using a small number of top-ranked features not only reduces model complexity but often leads to improved generalization.

While our study focuses on the prediction of the radius of gyration, we note the presence of consistent outliers across all architectures—such as OPN and ProtaN—which suggests opportunities for further refinement of model parameters or the incorporation of additional sequence-specific factors.. Finally, the current approach provides a foundation for extending the model to multi-output architectures that can capture a broader range of structural and dynamic properties of IDPs, including measures of heterogeneity and temporal evolution, thereby advancing our ability to characterize these complex biomolecules more comprehensively.

## VII. Acknowledgments

The research at UCF was supported by a DARPA AIBTO (HR00112530047) award. All computations were carried out using STOKES High-Performance Computing Cluster at UCF and through NSF-ACCESS grant (AB and SI) at the Purdue Anvil cluster.

