## Supplementary Materials for "Prediction of physical characteristics of disordered proteins using molecular simulation and physics-informed multiple machine learning strategies"

Diego Linares Gonzalez[1], Shahana Ibrahim[1,*], Swarnadeep Seth[2], George Atia[1,*], and Aniket Bhattacharya[2*]
[1]*Department of Electrical and Computer Engineering, University of Central Florida, Orlando, Florida 32816-2385, USA and*
[2]*Department of Physics, University of Central Florida, Orlando, Florida 32816-2385, USA*

### I. Input Parameters

In this section we provide details of the parameters introduced in Table-I in the main text. We define

$$M = \sum_{i=1}^{N} m_i, \quad \bar{m} = M/N \tag{S1a}$$

$$Q_{abs} = \sum_{i=1}^{N} |q_i|, \quad q_{abs} = Q_{abs}/N \tag{S1b}$$

$$Q_{net} = \sum_{i=1}^{N} q_i, \quad q_{net} = Q_{net}/N \tag{S1c}$$

$$\lambda_{sum} = \sum_{i=1}^{N} \lambda_i \quad \langle\lambda\rangle = \lambda_{sum}/N \tag{S1d}$$

Here, $m_i, q_i$, and $\lambda_i$ are the mass, charge, and hydropathy index of the $i^{th}$ amino acid. We include average mass per unit length $\bar{m}$ and average hydropathy $\langle\lambda\rangle$ in the training set of ANN and the quantities as itemized below.

Drawing analogy with the physical center of mass of a chain, we define three quantities $M_{CM}, Q_{CM}, \lambda_{CM}$, with respect to the center of the AA index $(N/2)$ as follows:

$$M_{CM} = \frac{1}{M}\sum_{i}^{N} \frac{i - i_{CM}}{i_{CM}} m_i, \tag{S2a}$$

$$Q_{CM} = \frac{1}{Q_{abs}}\sum_{i}^{N} \frac{i - i_{CM}}{i_{CM}} q_i. \tag{S2b}$$

$$\lambda_{CM} = \frac{1}{\lambda_{sum}}\sum_{i}^{N} \frac{i - i_{CM}}{i_{CM}} \lambda_i. \tag{S2c}$$

The value of $i_{CM}$ is the midpoint of $N$, either $N/2$ or $(N-1)/2$ depending upon the even and odd number of residues. Under this construction, the range $M_{CM}, Q_{CM}, \lambda_{CM}$ can vary between $(-1, 1)$ that captures the asymmetry in the residue mass, charge, and hydropathy distributions regardless of the IDP length.

Likewise, we define three RMS fluctuations in mass, charge, and hydropathy

$$\delta M = \sqrt{\frac{1}{N}\sum_{i}^{N}(m_i - \bar{m})^2} \tag{S3a}$$

$$\delta Q = \sqrt{\frac{1}{N}\sum_i^N (q_i - q_{abs})^2} \tag{S3b}$$

$$\delta\lambda = \sqrt{\frac{1}{N}\sum_i^N (\lambda_i - \langle\lambda\rangle)^2}. \tag{S3c}$$

Entropy in the sequence space plays an essential role to capture the sequence dependent properties of the IDPs. We calculate the dimensionless Shannon entropy $S_{IDP}^{all}$ and the Shannon entropy for the charged amino acids in the sequence $S_{IDP}^{q}$ (in units of Boltzmann constant $k_B$) of the FASTA sequence defined as follows

$$S_{IDP}^{all} = -\sum_i p_i \log_2(p_i). \tag{S4a}$$

$$S_{IDP}^{q} = -\sum_\alpha p_\alpha \log_2(p_\alpha). \tag{S4b}$$

Here $p_i = n_i/N$, where the $n_i$ is the number of times a particular amino acid with the index $i \in 20$ occurs in the sequence. Likewise, for $S_{IDP}^{q}$, $p_\alpha = n_\alpha/N_q$, and the index $\alpha$ runs for the charged amino acids only with $\sum_\alpha n_\alpha = N_q$. The repetitive appearance of a particular amino acid will reduce the number of permutation and its contribution in dynamical heterogeneity will be captured by the corresponding Shannon entropy.

The other relevant quantities used to train the ANN are as follows:

$$f_q^{+1} = N_q^{+1}/N; \quad f_q^{1/2} = N_q^{1/2}/N \tag{S5a}$$

$$f_q^{-1} = N_q^{-1}/N; \quad f_q^{0} = 1 - f_q^{+1} - f_q^{1/2} - f_q^{-1} \tag{S5b}$$

$$f^* = \frac{(f^+ - f^-)^2}{f^+ + f^-}, \tag{S5c}$$

$$SCD = \frac{1}{N}\sum_{i=2}^{N}\sum_{j=1}^{i-1} q_i q_j \cdot |i-j|^{0.5} \tag{S5d}$$

Here $N_q^{+1}$, $N_q^{1/2}$, and $N_q^{-1}/N$ are the number of IDPs with $+1$, $+1/2$, and $-1$ charges, so that $f_q^{+1}$, $f_q^{1/2}$, $f_q^{-1}$ are the corresponding fractions, and $f_q^0 = \left(N - N_q^{+1} - N_q^{+1/2} - N_q^{-1}\right)/N = N_q^0/N$ is the fraction of neutral amino acids. The quantities $f^+ = f_q^{+1} + f_q^{1/2}$ and $f^- = f_q^{-1}$ are the net positive and negative charge per residue of an IDP. The importance of charge asymmetry parameter $f^*$ [S1, S2] and the sequence charge decoration parameter $SCD$ parameter have been discussed in the literature [S3].

Finally, the quantities

$$l_{patch}^{(+1)}, \quad l_{patch}^{(+\frac{1}{2})}, \quad l_{patch}^{(-1)} \tag{S6}$$

are the contiguous charge patches consisting of unit positive, half-positive, and unit negative charges respectively (please note that the IDPs have $\pm1$ and $+\frac{1}{2}$ charges). $l_{patch}^{(q)} \neq 0$, for $q = \pm1, \frac{1}{2}$ if at least two consecutive amino acids have the same charge within the charge sequence (else $l_{patch}^{(q)} = 0$). We introduce the Sequence-Based Compactness Score (SBCS) metric to evaluate the compactness of the IDP based on the distribution and clustering of high-$\lambda$ residues. It is defined as

$$\text{SBCS} = \langle\lambda\rangle \sum_{j=1}^{k-1} \frac{1}{d_{j,j+1}} \tag{S7}$$

where, $d_{j,j+1}$ is the is the distance between consecutive high-$\lambda$ (hydrophobic) residues, indexed by $j$ and $j+1$, whose $\lambda > 0.5$ in the sequence, and $k$ is the total number those residues. The SBCS provides insights into how high-$\lambda$ residues are distributed within a sequence, and their potential role in influencing the aggregation behavior. It combines both the hydrophobicity of residues and their spatial organization to measure the overall compactness of the sequence.

The parameters as described in Eqns. (S1a) - (S7) along with the length $L$ of the amino acid constitute 23 important physics based features derived from the IDP sequence information as input vectors to train the MLP network. The distribution of the physical features are shown in Fig. 1 of the main text. We have investigated the relative weights these quantities through SHAP in Fig. S1 [S4, S5].

### II. SHAP Analysis

In Fig. S1, we present a SHAP analysis for each model that incorporates feature input. In the main text, we provided arguments that an approximate scaling law $\langle R_g \rangle = 0.38L^{0.56}$ is obeyed by the IDPs despite the fact that they are charged and have unequal masses, radii, and hydropathy interactions as demonstrated in our previous papers [S6, S7]. Thus, that this analysis shows that length remains the dominant variable across all hybrid models is not at all surprising. Beyond length, extracting the relative importance of other variables needs to be examined critically. Here we observe that depending on the architecture a few features stand out to be of larger relative weights as shown in Fig. S1. Since some of the features are not independent (for example the parameter $f^* = \frac{(f^+ - f^-)^2}{f^+ + f^-}$ [S2, S3] is dependent on the fraction of positive and negative charge per unit length - an architecture may give it a higher weight than the other. But this analysis with different architecture asserts that one can use a subset of the features to train an architecture and avoid overfitting issues. The integrated gradient analysis [S8] in the next section provide further relevant information.

### III. Integrated Gradient Analysis

In addition to SHAP analysis, the details of the *Integrated Gradients* analysis [S8] for each model is shown in Fig. S2. The analysis revealed that the top three most important features across all models were consistently Length, $f^*$, and SCD. Notably, SHD and the hydropathy parameter $\langle \lambda \rangle$ appear to be the 4th and 5th most important features in many evaluated models. Thus one can possibly obtain the same level of precision with these five parameters.

### IV. Attention Heatmaps for Interpretability

While one of the components of our method includes the assignment of attention weights to each input residue, we are able to plot those attention scores to gain insights into which regions of intrinsically disordered protein (IDP) sequences are most influential for the model's prediction. In this context, high attention weights indicate residues that are critical for the prediction task or are involved in highly important interactions. We can see some examples of attention maps in Figs. S4 and S5.

To further explore the interpretability of the model, we examined the distribution of attention scores across different amino acid types. Interestingly, the average attention weights did not vary dramatically across residues, suggesting that the model does not rely heavily on any single residue type to make its predictions.

These findings are consistent with the SHAP analysis results, which identified sequence length as the most influential feature, demonstrating that an increased number of residues corresponds to a larger radius of gyration.

Instead, the attention appears to be moderately distributed, with some focus on residues such as glutamic acid (E), alanine (A), glycine (G), proline (P), and asparagine (N). The overall uniformity across residues implies that the model is learning more complex, context-dependent representations rather than focusing on specific residues.

Moreover, to dive more into the residue type importance, we analyzed the average attention weights assigned to each residue, as illustrated in Figure S3. To mitigate bias introduced by varying sequence lengths , we first computed the average attention weight per residue type within each sequence. These per-sequence averages were then aggregated across the dataset, ensuring that longer sequences did not disproportionately influence the overall results.

---

[S1] Albert H Mao, Scott L Crick, Andreas Vitalis, Caitlin L Chicoine, and Rohit V Pappu. Net charge per residue modulates conformational ensembles of intrinsically disordered proteins. *Proceedings of the National Academy of Sciences*, 107(18):8183–8188, 2010.

[S2] Rahul K Das and Rohit V Pappu. Conformations of intrinsically disordered proteins are influenced by linear sequence distributions of oppositely charged residues. *Proceedings of the National Academy of Sciences*, 110(33):13392–13397, 2013.

[S3] Kingshuk Ghosh, Jonathan Huihui, Michael Phillips, and Austin Haider. Rules of physical mathematics govern intrinsically disordered proteins. *Annual Review of Biophysics*, 51(1):355–376, 2022.

[S4] Lloyd S Shapley et al. A value for n-person games. 1953.

[S5] Scott M Lundberg and Su-In Lee. A unified approach to interpreting model predictions. *Advances in neural information processing systems*, 30, 2017.

[S6] Jacob Bair, Swarnadeep Seth, and Aniket Bhattacharya. Universality in conformations and transverse fluctuations of a semi-flexible polymer in a crowded environment. *The Journal of Chemical Physics*, 158(20), 2023.

[S7] Swarnadeep Seth and Aniket Bhattacharya. Accelerated missense mutation identification in intrinsically disordered proteins using deep learning. *Biomacromolecules*, 26(4):2106–2115, 2025.

[S8] Mukund Sundararajan, Ankur Taly, and Qiqi Yan. Axiomatic attribution for deep networks, 2017.

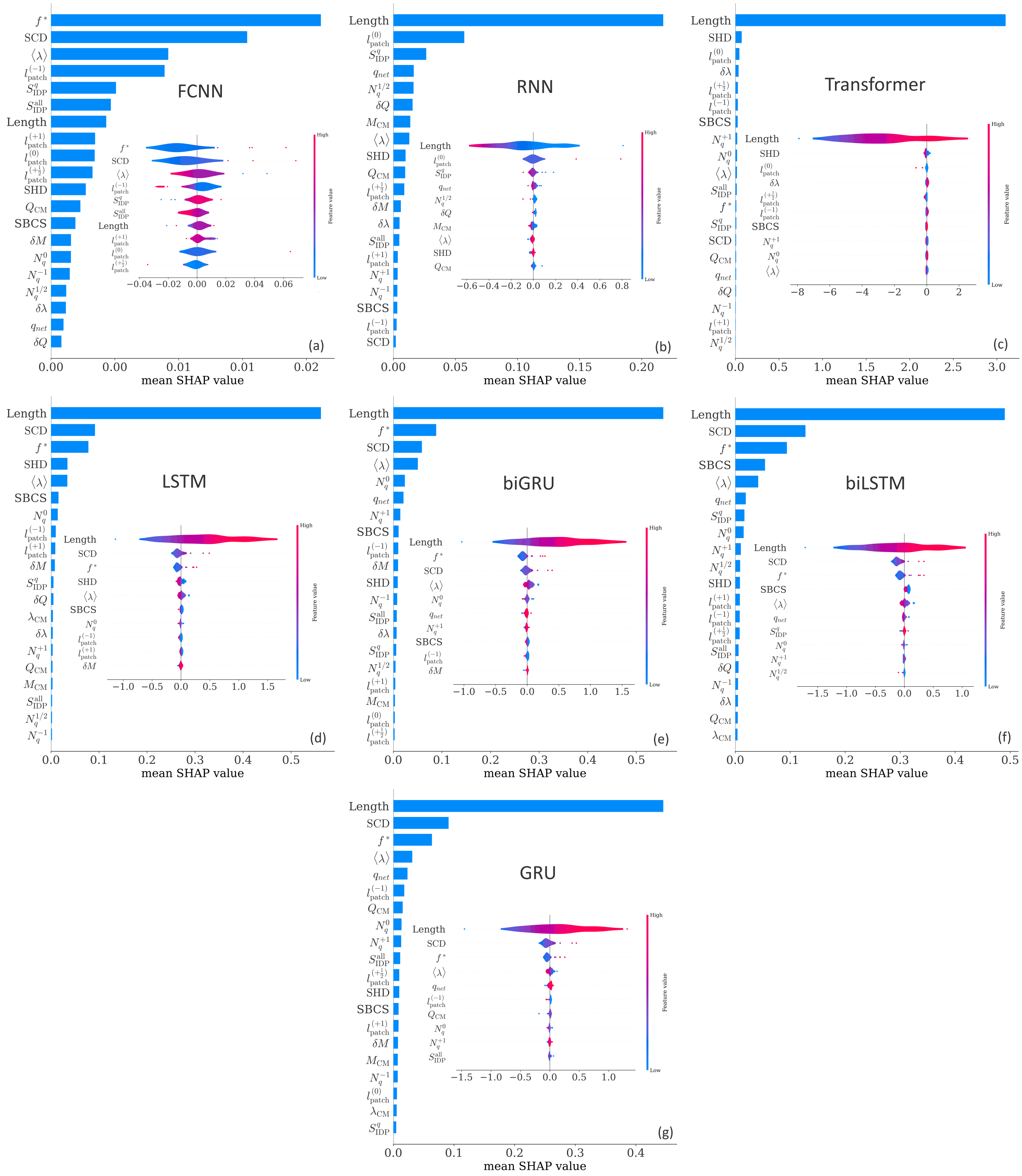


FIG. S1. SHAP feature importance summary for FCNN, RNN, Transformer, LSTM,biLSTM, GRU, biGRU

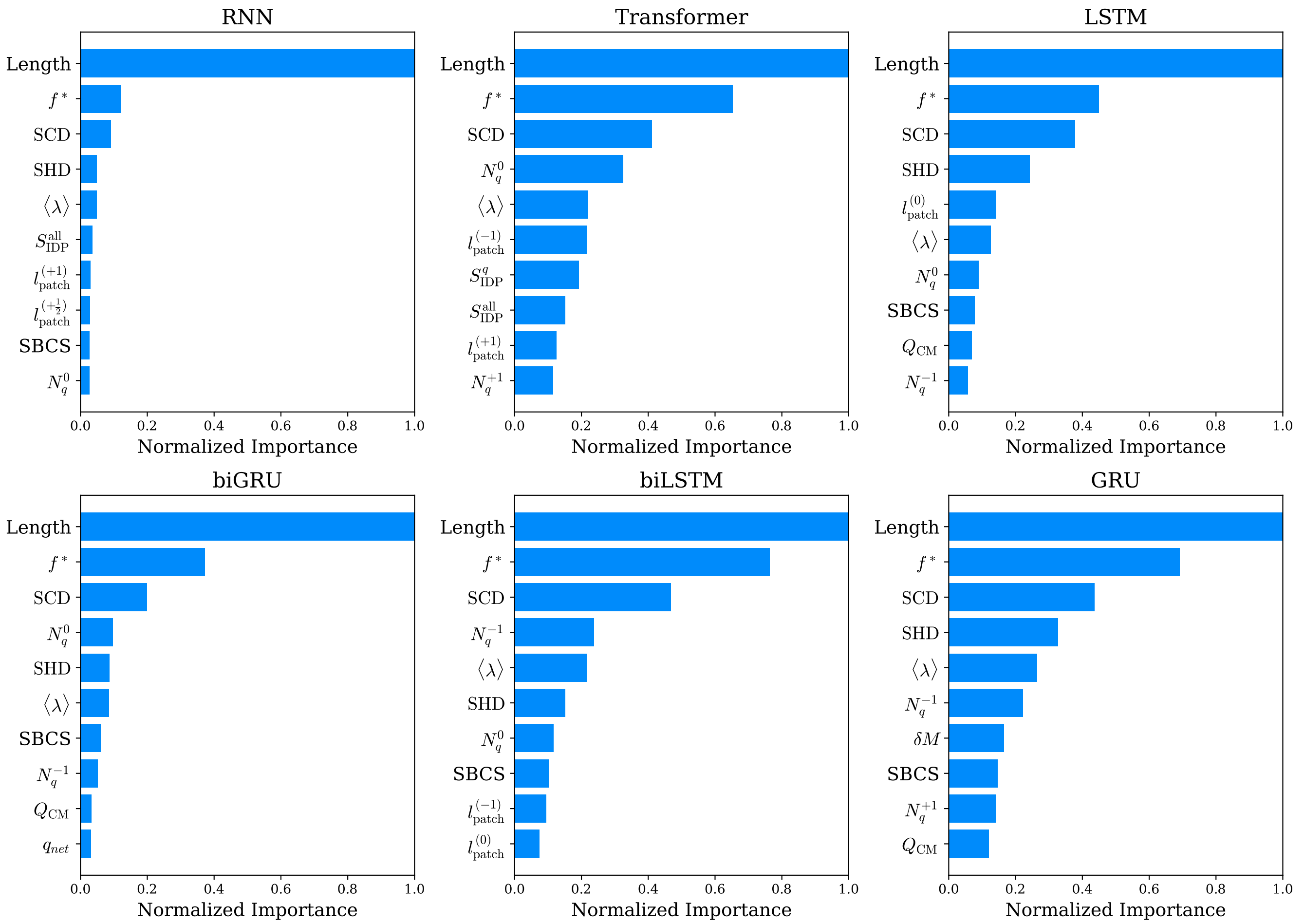


FIG. S2. Top 10 most important features identified using Integrated Gradients for each architecture. Feature importance is normalized.

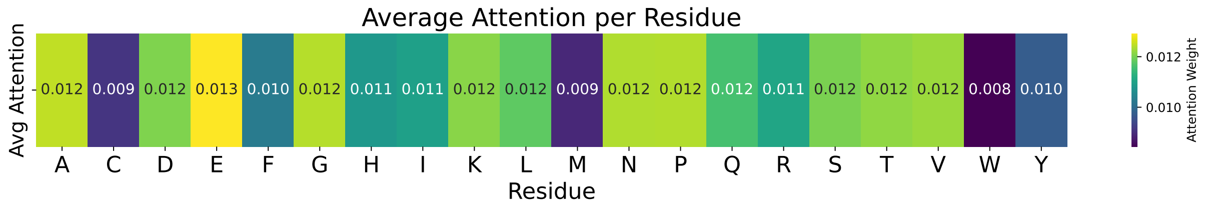


FIG. S3. Per-residue average attention weights and standard deviations across sequences, computed using per-sequence averaging with the Hybrid biGRU model.

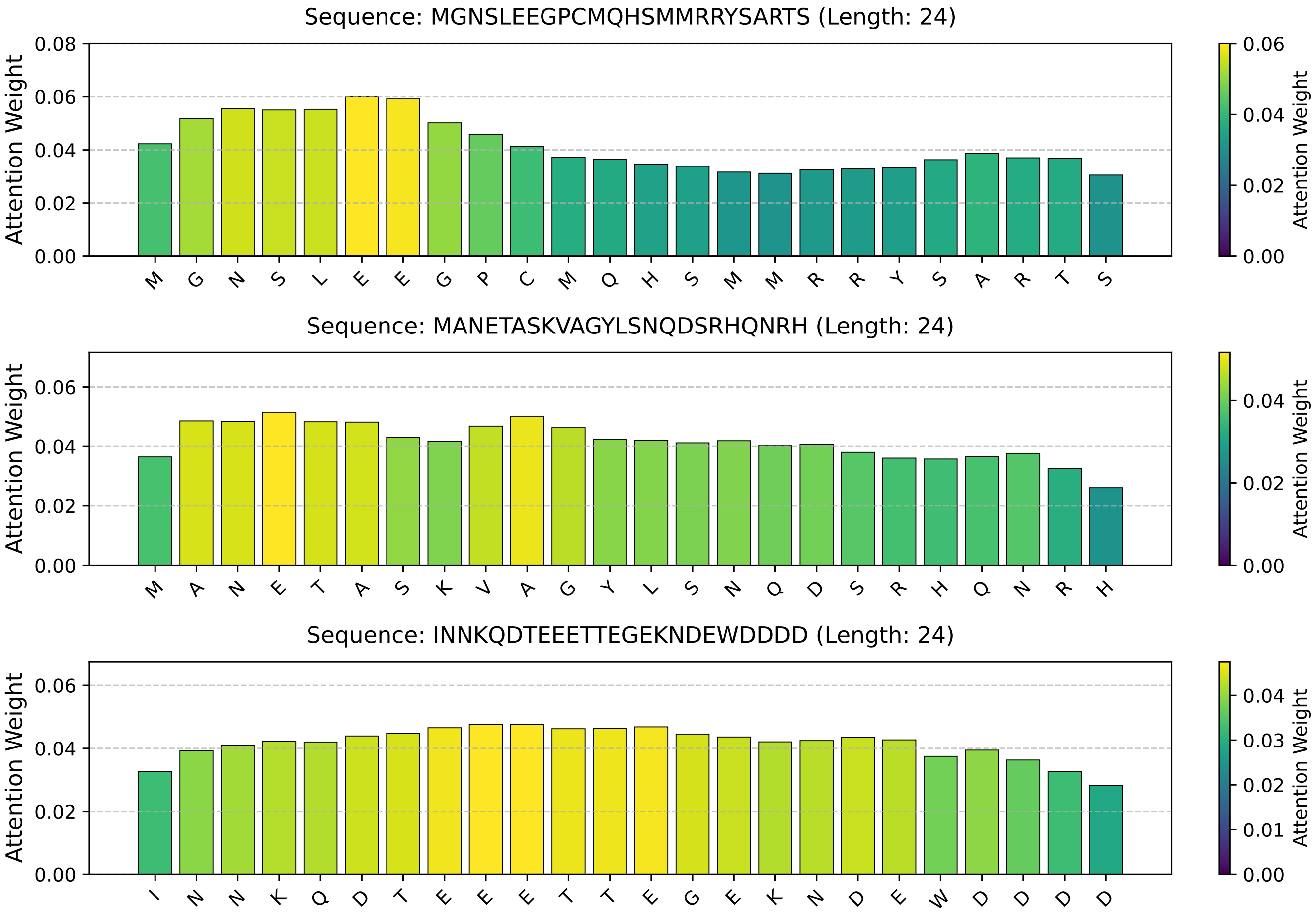


FIG. S4. Attention maps from the Hybrid biGRU model for three randomly selected sequences, each of length 24.

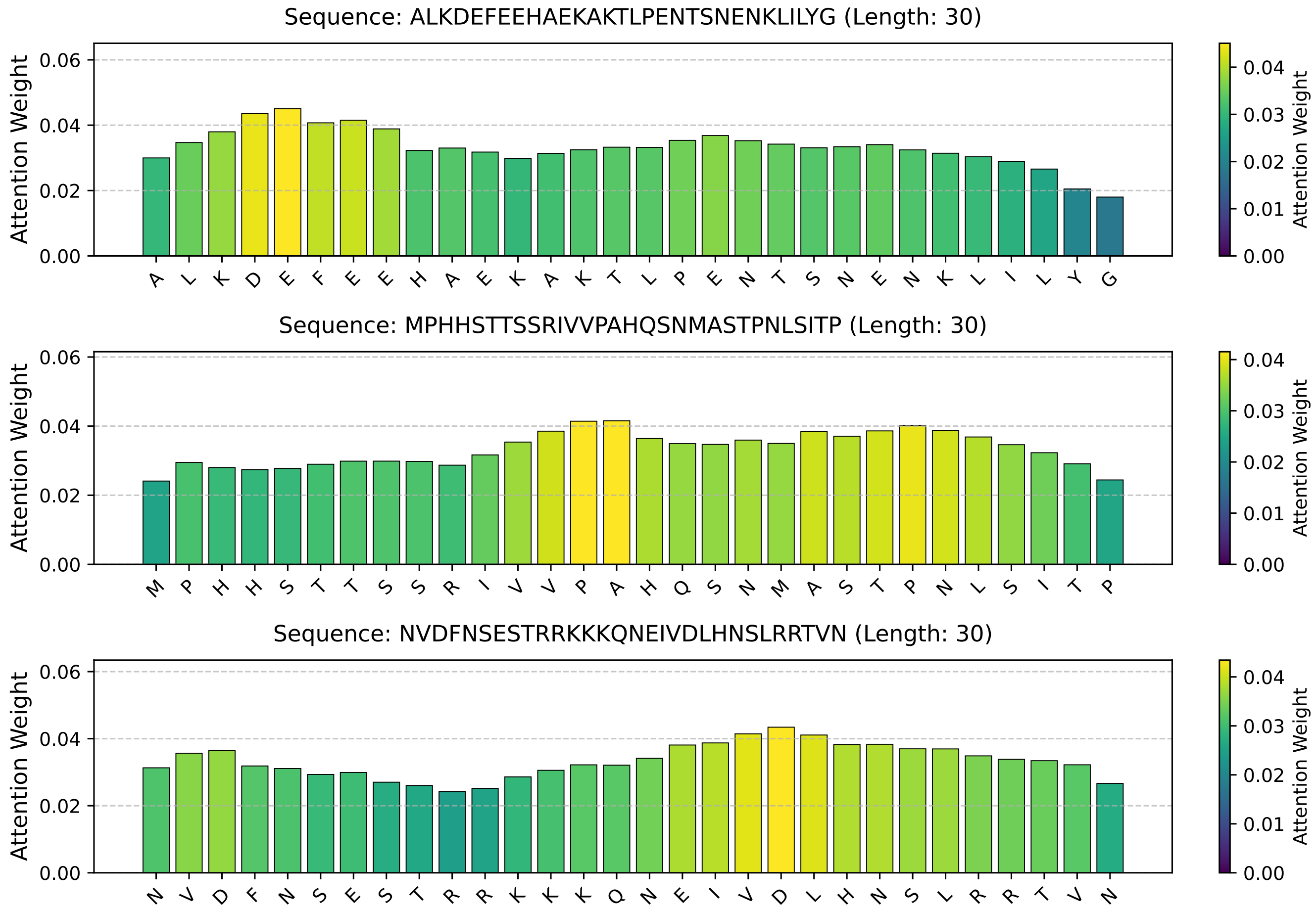


FIG. S5. Attention maps from the Hybrid biGRU model for three randomly selected sequences, each of length 30.
